# Dissociable roles of central striatum and anterior lateral motor area in initiating and sustaining naturalistic behavior

**DOI:** 10.1101/2020.01.08.899070

**Authors:** Victoria L. Corbit, Sean C. Piantadosi, Jesse Wood, Grace Liu, Clare J.Y. Choi, Ilana B. Witten, Aryn H. Gittis, Susanne E. Ahmari

## Abstract

Although much is known about how corticostriatal circuits mediate behavioral selection, most previous work has been conducted in highly trained animals engaged in instrumental tasks. Understanding how corticostriatal circuits mediate behavioral selection and initiation in a naturalistic setting is critical to understanding how the brain chooses and executes behavior in unconstrained situations. Central striatum (CS), an understudied region that lies in the middle of the motor-limbic topography, is well-poised to play an important role in these processes since its main cortical inputs (Corbit et al., 2019) have been implicated in behavioral flexibility (lateral orbitofrontal cortex (Kim and Ragozzino, 2005)) and response preparation (anterior lateral motor area, ALM) (Li et al., 2015), However, although CS activity has been associated with conditioned grooming behavior in transgenic mice (Burguiere et al., 2013), the role of CS and its cortical inputs in the selection of spontaneous behaviors has not been explored. We therefore studied the role of CS corticostriatal circuits in behavioral selection in an open field context.

Surprisingly, using fiber photometry in this unconstrained environment, we found that population calcium activity in CS was specifically increased at onset of grooming, and not at onset of other spontaneous behaviors such as rearing or locomotion. Supporting a potential selective role for CS in the initiation of grooming, bilateral optogenetic stimulation of CS evoked immediate onset grooming-related movements. However, these movements resembled subcomponents of grooming behavior and not full-fledged grooming bouts, suggesting that additional input(s) are required to appropriately sequence and sustain this complex motor behavior once initiated. Consistent with this idea, optogenetic stimulation of CS inputs from ALM generated sustained grooming responses that evolved on a time-course paralleling CS activation monitored using single-cell calcium imaging. Furthermore, fiber photometry in ALM demonstrated a gradual ramp in calcium activity that peaked at time of grooming termination, supporting a potential role for ALM in encoding length of this spontaneous sequenced behavior. Finally, dual color dual region fiber photometry indicated that CS activation precedes ALM during naturalistic grooming sequences. Taken together, these data support a novel model in which CS activity is sufficient to initiate grooming behavior, but ALM activity is necessary to sustain and encode the length of grooming bouts. Thus, the use of an unconstrained behavioral paradigm has allowed us to uncover surprising roles for CS and ALM in the initiation and maintenance of spontaneous sequenced behaviors.

## Introduction

In order to navigate our world, it is essential to string together series of movements into complete actions that can be used to attain goals. Carrying out actions in the appropriate context and for the appropriate amount of time is essential for adaptive behavior. However, although much information has been gained recently regarding the neural substrates underlying these processes (Balleine and O’Doherty, 2010; Cui et al., 2013; Frank, 2011; Jin et al., 2014; Tecuapetla et al., 2016), the mechanisms underlying both appropriate initiation and sustainment of spontaneous actions are still incompletely understood.

Prior work has implicated striatal circuits in behavioral selection. In highly trained animals, it has been shown that dorsal striatal spiny projection neurons (SPNs) have activity that correlates with the first and/or last action in a sequence of movements, suggesting that striatum encodes movement sequences as chunked actions (Graybiel, 1998; Jin et al., 2014; Tecuapetla et al., 2016). Furthermore, lesions of the lateral or medial regions of dorsal striatum can cause a well-trained animal to exhibit goal-directed or habitual behavior, respectively, on a lever press task (Yin et al., 2004; Yin et al., 2005). These data thus suggest that specific regions of dorsal striatum and/or their upstream inputs are important for modulating the extent to which behavior is intentional (e.g. value-based) or automatic (e.g. value-independent).

While these operant-based paradigms have yielded great insight into the role of the striatum in learning and performing trained actions, observing naturalistic behavior is a better approximation for understanding the neural mechanisms of key behaviors essential for survival. For instance, mice spontaneously engage in rearing and locomotor behavior to explore their surroundings; these behaviors are essential for obtaining food and avoiding threats in the environment. In addition, rodents perform behaviors such as grooming and nest-building to maintain hygiene and care for pups. Unfortunately, despite the importance of these behaviors for rodent survival, only a few studies have investigated the role of the striatum in behavioral selection in unrestrained, untrained settings. Using both fiber photometry (Cui et al., 2013; Markowitz et al., 2018) and one-photon miniature microscope calcium imaging (Barbera et al., 2016; Parker et al., 2018), several of these studies have demonstrated activation of dorsal striatal SPNs prior to or at the onset of locomotion (Barbera et al., 2016; Cui et al., 2013; Markowitz et al., 2018; Parker et al., 2018), turning (Cui et al., 2013; Markowitz et al., 2018; Parker et al., 2018), and rearing (Markowitz et al., 2018; Parker et al., 2018). These results suggest that diverse types of spontaneous movements are associated with increased activity in dorsal striatum, though the specific SPN temporal activation profiles for different behaviors in relation to movement onset demonstrate some variation. However, because these studies were associational, and prior causal studies focused solely on the role of dorsal striatum in locomotion (Kravitz et al., 2010), further work is needed to determine whether particular striatal activity patterns can directly generate species-typical behaviors.

The striatum exhibits topographical cortical inputs following a motor-limbic gradient in the dorsal to ventral direction (Alexander et al., 1986; Haber, 2016). Interestingly, this positions central striatum (CS) at a crucial nexus of corticostriatal inputs along this topography (Ebrahimi et al., 1992; Oh et al., 2014), making it well suited to link motivational factors and sensorimotor responses. Consistent with this idea, in prior work we demonstrated that CS receives projections from both lateral orbitofrontal cortex (LOFC) and anterior lateral motor area (ALM) (Corbit 2019), regions implicated in behavioral flexibility (Kim and Ragozzino, 2005) and motor preparation (Guo et al., 2014; Li et al., 2015), respectively. Furthermore, early studies showed that activating central regions of striatum via disinhibition with picrotoxin caused tic-like behavior in rodents (Tarsy et al., 1978). More recent work has replicated this picrotoxin effect and further showed that the production of tics is dependent on CS activity, and not cortical activity (Pogorelov et al., 2015). The production of tics by disinhibition of CS highlights a potential role for this region in initiation of movement fragments.

Despite this convergent evidence that hints at an important role for CS in selection of spontaneous behavior, little has been done to investigate this hypothesis in vivo. One prior study showed CS hyperactivity at baseline and during persistent conditioned grooming behavior in the SAPAP3-KO transgenic mouse model of compulsive behavior (Burguiere et al., 2013), suggesting a correlation between abnormal activity in CS and abnormal behavioral selection. Burguiere and colleagues also showed that activating LOFC inputs to CS was sufficient to reduce this grooming behavior, suggesting that LOFC serves as an inhibitor of abnormal behavior generated in CS (Burguiere et al., 2013). However, CS inputs from ALM, a region that has been associated with preparation, sustainment, and sequencing of trained behavior (Guo et al., 2014; Li et al., 2015; Rothwell et al., 2015; Xu et al., 2019), have not yet been examined. We therefore sought to determine whether CS and its inputs from ALM mediate spontaneous naturalistic behavior.

To investigate the role of ALM-CS corticostriatal circuits in naturalistic behavioral selection, we used fiber photometry, optogenetics, and *in vivo* microendoscopy to observe and manipulate these circuits in vivo. Surprisingly, using fiber photometry, we found a selective increase in CS activity at the initiation of grooming events but not at onset of other spontaneous behaviors. Direct optogenetic stimulation of CS SPNs to mimic this increase in activity evoked immediate-onset grooming-related fragmented movements that were short in duration. Using fiber photometry, we then observed that ALM is also selectively activated during grooming behavior. Surprisingly, peak ALM activity correlated with grooming bout length, suggesting that ALM encodes duration of sequenced behaviors. In contrast to CS direct stimulation, stimulation of ALM terminals in CS caused long-latency bouts that resembled normal grooming behaviors. Finally, dual-color, dual-region photometry revealed that CS activation precedes ALM activation at the initiation of naturalistic grooming bouts. Taken together, these results suggest that CS selects or initiates naturalistic grooming behavior, while ALM activity plays a role in sustaining and properly compiling grooming-related movements into effective bouts.

## Results

### Increased activity in CS is specifically associated with groom start

To investigate the role of CS activity in naturalistic behaviors, we unilaterally injected AAV-JRGECO1a into CS and implanted a 200um fiber photometry probe 300um above the injection site (Fig.1A). Bulk calcium activity was recorded while animals performed unconstrained, spontaneous behaviors in an observation chamber. CS calcium activity was then analyzed at onset of 3 distinct spontaneous behaviors that comprise a majority of non-immobility time: grooming, rearing, and locomotion (Fig.1B). Strikingly, we observed that CS showed a rapid increase in activity at the onset of grooming (Fig.1C, repeated-measures ANOVA, time x virus interaction p = .03). To further delineate which aspect of grooming is encoded by CS, we separated bouts into face grooming (which primarily involves paw movements) or body grooming (which primarily involves tongue and body movements). This analysis demonstrated that increased CS activity at grooming onset is primarily associated with body grooming (Fig.1D, repeated-measures ANOVA, time x behavior type interaction p = .002). In contrast, when we aligned activity to the onset of rearing, we did not observe increased CS activity, instead seeing a significant reduction in activity after rear onset (repeated-measures ANOVA, time x virus interaction p <.001, Fig.1E). Similarly, CS activity did not increase at locomotion onset (repeated-measures ANOVA, time x virus interaction p = .55, Fig.1F). Taken together, these data suggest that CS activity may be specialized for the generation of sub-components of grooming behavior related to body grooming, such as licking the flank or torso, but do not prove a causal role for CS in their generation.

**Figure 1.**
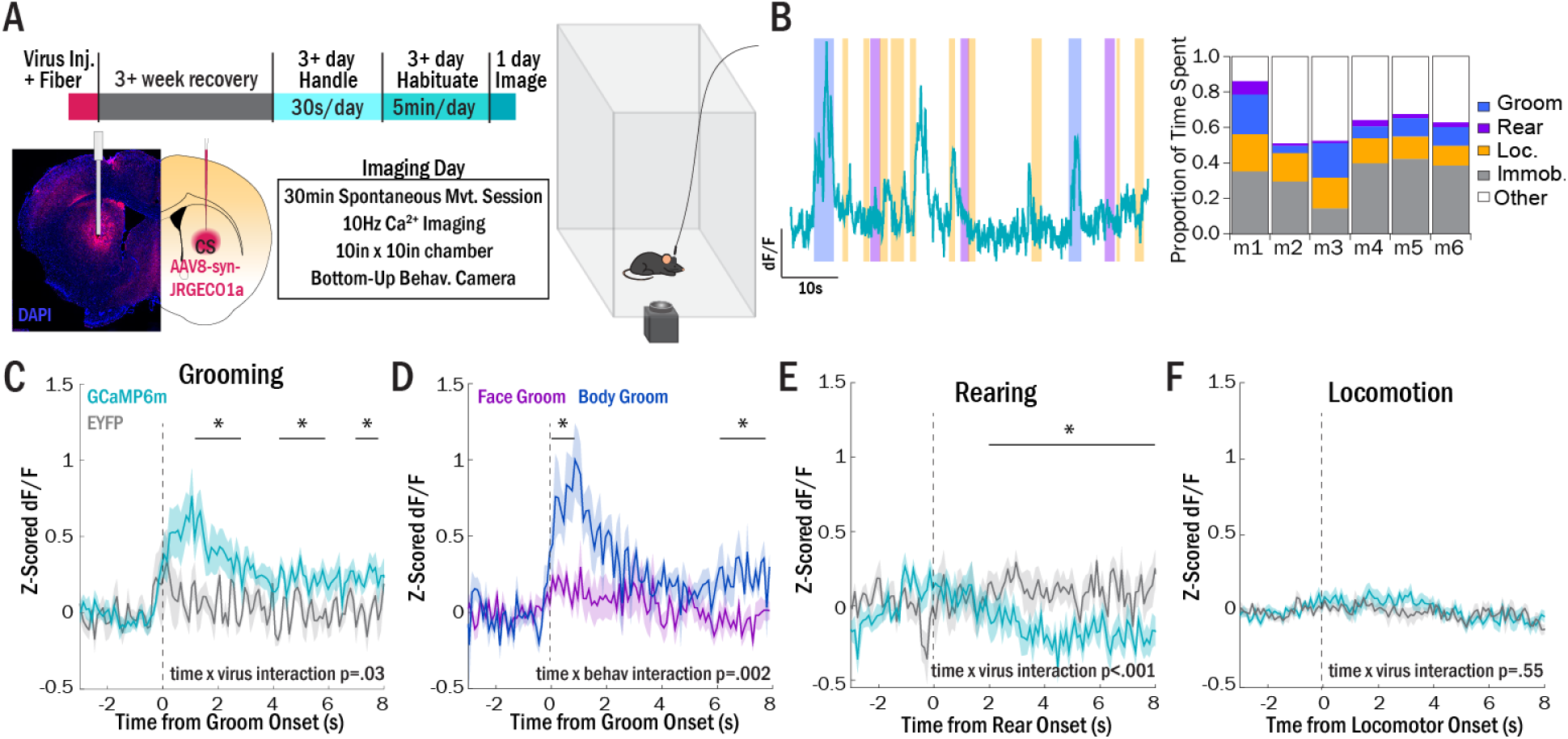
CS calcium activity increases selectively at the onset of grooming behavior. (A) Mice were injected with AAV8-syn-JRGECO1a unilaterally in central striatum (CS). After at least 3 weeks recovery, mice were handled for at least 3 days for 30s/day, and then habituated for at least 3 days to scruffing, cable attachment, and the observation chamber. Following habituation, mice were placed in a bottom-up observation chamber and CS bulk calcium activity was recorded using fiber photometry for 30 minutes during spontaneous behavior. (B) (left) Example trace of CS calcium activity aligned to 3 distinct behaviors: grooming, rearing, and locomotion. (right) Proportions of each subclass of behavior in 6 example mice. “Other” includes combinations of sniffing, fine head and paw movements, and postural adjustments. (C) Normalized CS calcium activity aligned to onset of grooming behavior (GCaMP6m: N=9 (teal); GFP control: N=5 (gray), two-way repeated measures ANOVA, time x virus interaction: p = .03, significant post-hoc contrasts (p < .05) for bins 1-3s, 4-6s, and 7-8s relative to baseline (1s before onset)). (D) Normalized GCaMP6m activity for grooming subtypes: face grooming (purple); body grooming (blue) (two-way repeated measures ANOVA, time x behavior interaction: p = .002, significant post-hoc contrasts (p < .05) for bins 0-1s and 6-8s relative to baseline (1s before onset)). (E) Normalized CS calcium activity aligned to onset of rearing behavior (GCaMP6m: N=9 (teal); GFP control: N=5 (gray), two-way repeated measures ANOVA, time x virus interaction: p < .001, significant post-hoc contrasts (p < .05) for bins 2-8s relative to baseline (1s before onset)). (F) Normalized CS calcium activity aligned to onset of locomotion (GCaMP6m: N=9 (teal); GFP control: N=5 (gray), two-way repeated measures ANOVA, time x virus interaction: p = .55).

### Optogenetic stimulation of CS evokes partial grooming movements

To test whether increased activity in CS could directly cause grooming behavior, we bilaterally injected hSyn-ChR2-EYFP or EYFP control virus into the CS of WT mice and implanted optical fibers above the injection site (Fig.2A). Because our photometry data suggested that grooming-related CS activity entails an increase from baseline, we developed a stimulation paradigm that enabled us to capture a sufficient number of trials during which animals were quiescent and therefore presumably at baseline CS activity levels [20s constant light pulses; pseudorandom inter-trial interval (30s+/-5s)] (Fig.2B).

**Figure 2.**
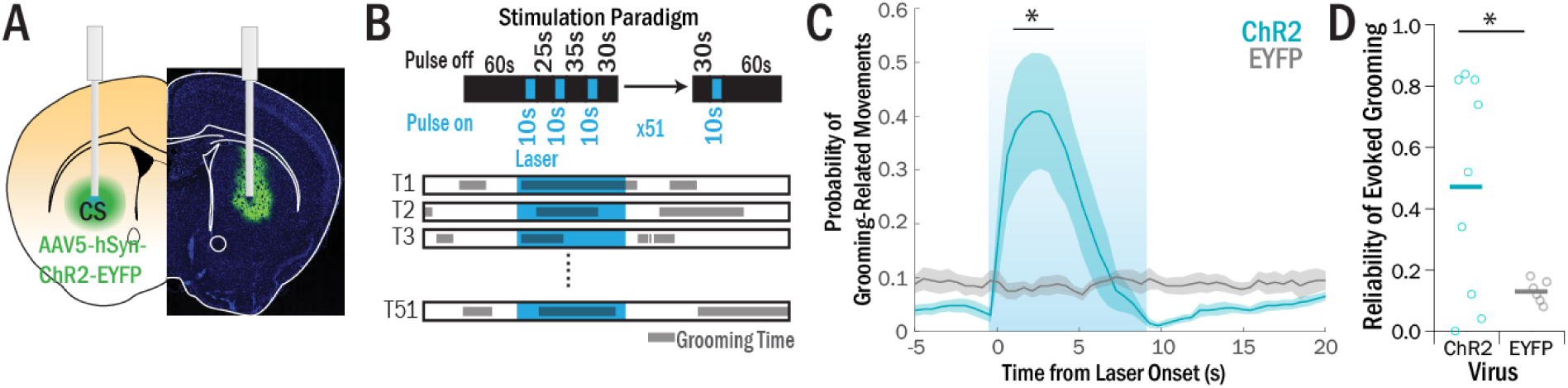
CS optogenetic stimulation evokes grooming-like movement fragments with short latency. (A) AAV5-syn-ChR2-EYFP was bilaterally injected in CS and fiberoptics were implanted above each injection site. (B) 51 stimulation trials of 10s constant light were separated by pseudorandom inter-trial interval of 25-35s with a 5s jitter. Grooming and grooming-related movement fragments were manually scored and aligned to each laser presentation, and peri-laser grooming probability was calculated. (C) Probability of grooming or grooming-related movements in ChR2 mice (N=9) was significantly greater than in EYFP mice (N=6) at the onset of the laser pulse (two-way repeated measures ANOVA, significant time x virus interaction, significant in bins 1-3.5s after laser onset). (D) Reliability of evoking a grooming response was calculated by dividing the number of trials in which an animal started grooming or performing grooming movement fragments during laser-on time by the total number of trials (51). ChR2 mice had significantly greater reliability of an evoked response (47%) relative to EYFP mice (0.13%) (t-test, p=.01).

The primary response to CS stimulation was the initiation of grooming like movements (Fig.2C, Supp. Video 1)-that is, evoked behavior resembled grooming behavior but appeared fragmented. In addition, grooming-like movements were stereotyped–i.e., light pulses consistently evoked qualitatively similar movements within a given animal. To quantify the temporal dynamics of these laser-evoked responses, we calculated when grooming or grooming-like behaviors were evoked during the laser-on period. Laser-evoked grooming movements were initiated with a short latency after CS stimulation, averaging 1.03s (SEM=0.95s, Fig.2E). As a proxy for reliability of stimulation-evoked behavior within a mouse, we also assessed grooming probability as *# stim trials with evoked behavior/ # total stim trials*. ChR2 mice had a significantly greater reliability of laser-evoked grooming (44.0±.1%) than EYFP control mice (0.13±.04%, t-test, p = .04, Fig.2D) Interestingly, this effect appeared to be specific to grooming-like movements, as we did not see a similar increase in rearing or locomotion during CS stimulation (SuppFig.1A-B). Taken together, these data demonstrate that activation of CS selectively evokes short-latency “syllables” of grooming behavior. However, the absence of complete bouts of grooming behavior after CS stimulation suggested that an alternative brain region is necessary for appropriate sequencing of grooming-related movements into sustained bouts.

### ALM shows increased activity specifically at grooming onset

Prior work has demonstrated that anterior M2/anterior lateral motor area (ALM), one of the major cortical inputs to CS (Corbit et al., 2019), is associated both with appropriate sequencing of movements during trained tasks and with generating licking movements (Bollu et al., 2019; Li et al., 2015; Rothwell et al., 2015). ALM may therefore be uniquely suited for guiding selection of spontaneously generated behaviors such as body grooming. To investigate the relationship between ALM activity and the generation of untrained behaviors, we therefore performed unilateral fiber photometry in ALM to record calcium activity during grooming, rearing, and locomotion bouts in spontaneously behaving mice (Fig.3A-B).

**Figure 3.**
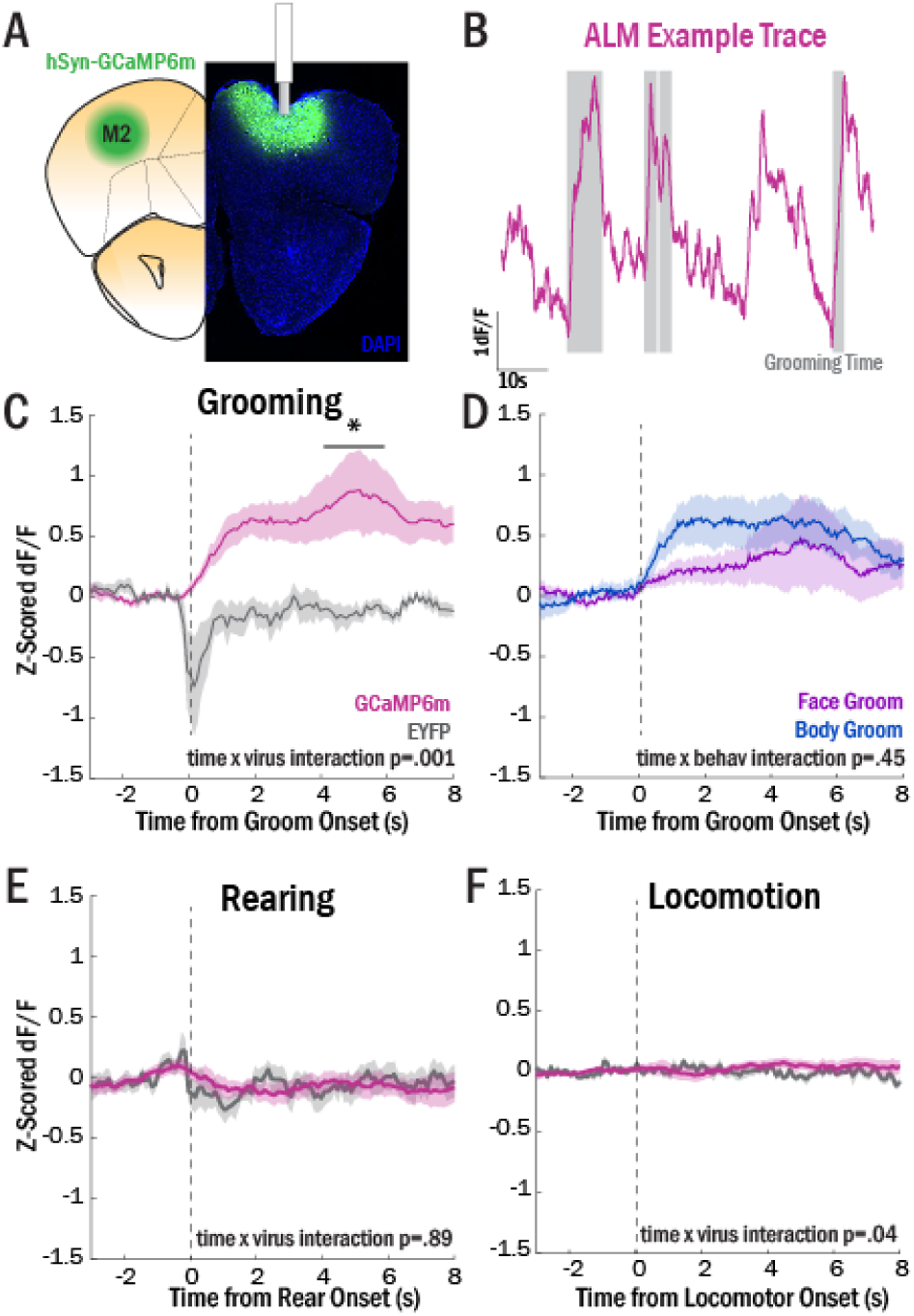
ALM calcium activity increases selectively at the onset of grooming behavior. (A) Fiber photometry experiments were conducted by injecting AAV9-GCaMP6m unilaterally into ALM and implanting a fiber optic 0.10mm over the injection. Habituation performed as in Fig.1A. (B) Example trace of ALM bulk calcium activity with grooming events highlighted in gray. (C) Normalized ALM calcium activity aligned to the onset of grooming behavior (GCaMP6m: N=9 (maroon); GFP control: N=6 (gray); two-way repeated measures ANOVA, time x virus interaction: p = .001, significant post-hoc contrasts (p < .05) for bins 4-6s relative to baseline (1s before onset). (D) Normalized GCaMP6m activity for grooming subtypes: face grooming (purple); body grooming (blue) (two-way repeated measures ANOVA, time x behavior interaction: p = .45). (E) Normalized ALM calcium activity aligned to onset of rearing behavior (GCaMP6m: N=9 (maroon); GFP control: N=6 (gray); two-way repeated measures ANOVA, time x virus interaction: p = .89). (F) Normalized ALM calcium activity aligned to onset of locomotion (GCaMP6m: N=9 (maroon); GFP control: N=6 (gray), two-way repeated measures ANOVA, time x virus interaction: p = .051)

In contrast to the proposed role of ALM and supplementary motor cortices in the preparation of actions (Lee et al., 1999; Li et al., 2015; Romo and Schultz, 1992), we found that ALM activity increased gradually at the onset of, and not prior to, grooming behavior (Fig.3C; repeated measures ANOVA, time x virus interaction p < .001). When we separated calcium traces based on grooming subtype (face vs. body), we observed a trend towards sharper increases in activity after body grooming onset (face slope = 0.06 ± 0.04, body slope = 0.15 ± 0.05, t(7)=-2.22, p=.06), but not overall significant differences between the traces (Fig.3D; repeated measures ANOVA, time x groom subtype interaction p =.45; time main effect p<.001). Similar to our observations in CS, we did not see increases in calcium activity at rearing onset (Fig.3E; repeated measures ANOVA, time x virus interaction p=.89) or at the onset of locomotion bouts (Fig.3F; repeated measures ANOVA, time x virus interaction p=.051).

### Peaks in ALM activity correlate with grooming cessation

Close examination of the temporal dynamics of the grooming-related calcium activity in ALM revealed that the average peak in fluorescence occurred several seconds after grooming onset (3.21 ± 2.77s post onset). This long latency to peak fluorescence suggested that, contrary to predictions, ALM activity is related to termination of a grooming bout, not initiation. To investigate this hypothesis more rigorously, we first inspected the ALM photometry data on a trial-by-trial basis, comparing calcium activity on a given grooming trial with the timing of initiation and termination of grooming behavior (Fig.4A). Interestingly, we observed that ALM activity typically increased at the start of a grooming bout and remained elevated until that bout was terminated (example traces in Fig.4B). To further probe this phenomenon, we separated grooming trials into quartiles (2s intervals) and calculated the average ALM calcium activity traces from each quartile (Fig.4C). This analysis revealed a pattern of longer time to peak fluorescence and increased peak height as grooming bout duration increased (Fig.4C). To better quantify this phenomenon, we identified the first prominent peak after groom onset in each trial and measured the time-to-peak and amplitude (see Methods for full description of quantification). In support of our findings from the analysis of grooming bout quartiles, we observed a significant correlation of bout length with ALM fluorescence peak time, suggesting that ALM grooming-related activity peaks at groom offset (Fig.4D, R^2^ = .40, p = 1×10^−14^). We also observed a weaker, but significant, correlation of bout length with peak amplitude (Fig.4E, R^2^=.13, p=4.8×10^−5^), suggesting that both the peak time and amplitude of ALM activity are good predictors of bout length. However, a multiple linear regression run on these two factors indicated that the predictive power of peak amplitude and peak time combined was only slightly greater than the predictive power of peak time alone (R^2^ = .41, p = 1.3×10^−14^). By contrast, a similar correlation between grooming bout length and CS calcium activity was not observed (SuppFig.2).

**Figure 4.**
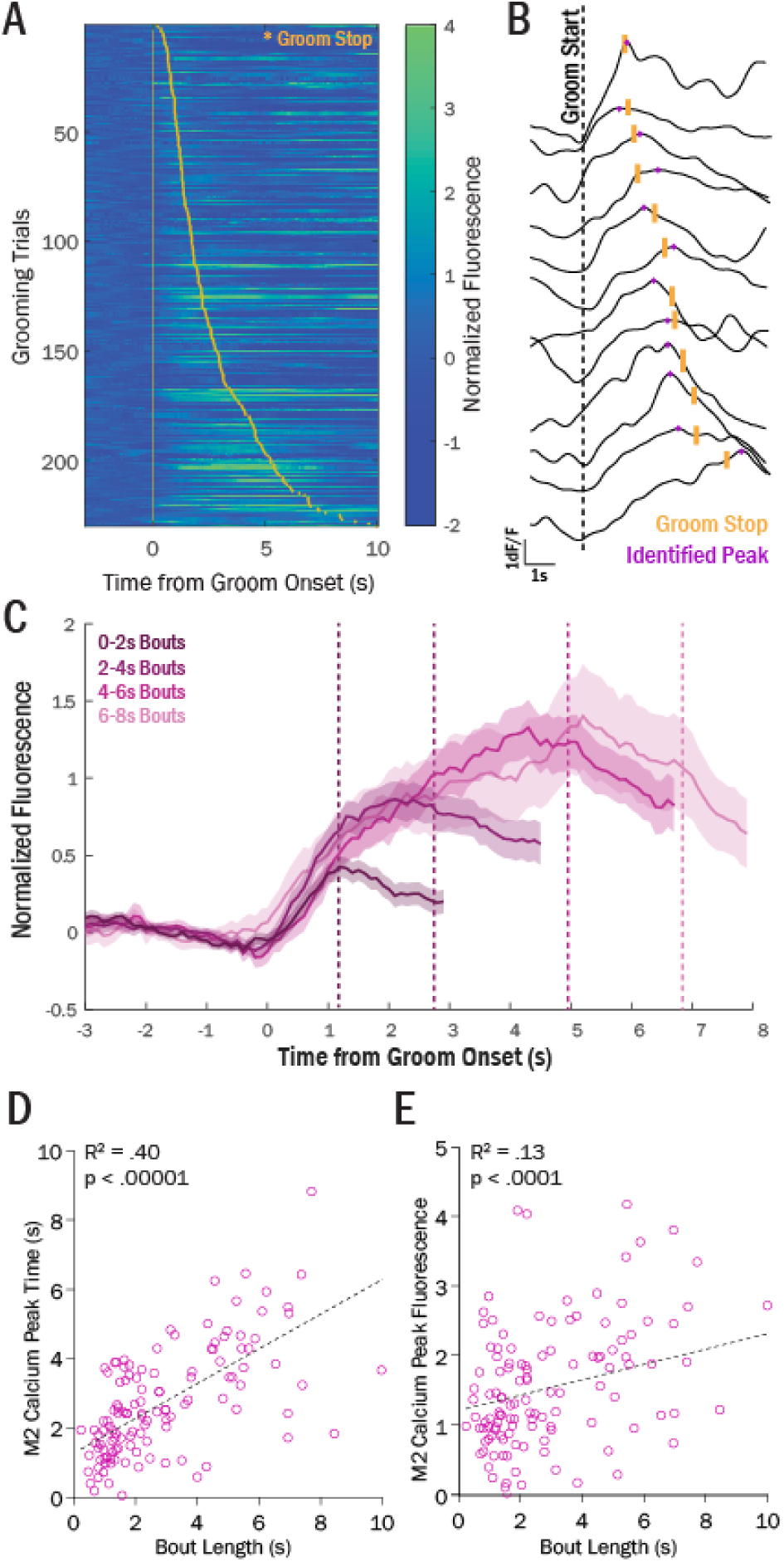
ALM calcium activity correlates with grooming bout length. (A) Heat map of ALM normalized fluorescence for individual trials of grooming behavior (baselined to -3:0s prior to groom onset). Grooming offset for each trial is marked with a yellow star. (B) Individual representative traces of ALM calcium traces show ramping activity between groom onset (dashed line) and groom offset (yellow hash). Peaks were identified using MATLAB’s “prominence” measurement (see Methods) – automatically identified peaks for each trial are labeled in purple. (C) Separation of grooming bouts into 4 quartiles (0-2s [N = 103 bouts], 2-4s [N = 53 bouts], 4-6s [N = 29 bouts], and 6-8s duration [N = 10 bouts]) reveals pattern of increasing peak amplitude and peak time in longer grooming bouts. Each dotted line marks average bout length for corresponding quartile (denoted by different shades of pink). Bouts > 8s were excluded in plot because of insufficient N (<5 bouts for each remaining quartile) (D) Time of detected peak significantly correlates with bout duration on trial-by-trial basis (Pearson correlation, R^2^ = .53, p 5.7×10^−23^). (E) Amplitude of detected peak significantly correlates with bout duration on trial-by-trial basis (Pearson correlation, R^2^ = .18, p = 2.0×10^−7^). For (D) and (E), X-axis cropped at 10s for clarity, though bouts > 10s were included in analysis (see SuppFig.2A).

Taken together, these photometry data suggest that ALM activity is preferentially associated with grooming behavior compared to other spontaneous behaviors. Furthermore, time to peak activity in ALM correlates with the length of naturally occurring grooming bouts, suggesting that ALM activity may play a role in sustaining grooming bouts, but is not directly responsible for initiating grooming behavior.

### ALM-CS stimulation evokes full grooming behavior at long latency

These combined data indicate that CS activity is related to grooming initiation, while ALM activity encodes grooming bout length, suggesting that ALM may be responsible for sequencing and sustaining fragments of grooming-related movements that are generated via activation of CS. According to this model, activating ALM terminals in CS should initiate smooth and sustained grooming bouts, in contrast to the fragmented movements generated by direct activation of CS. To test this hypothesis, we bilaterally injected ChR2-EYFP (or EYFP control virus) into ALM, and implanted optical fibers over ALM axon terminals in CS (Fig.5A). Terminal stimulation was performed with a pulsed stimulation paradigm (see Methods and SuppFig.3).

**Figure 5.**
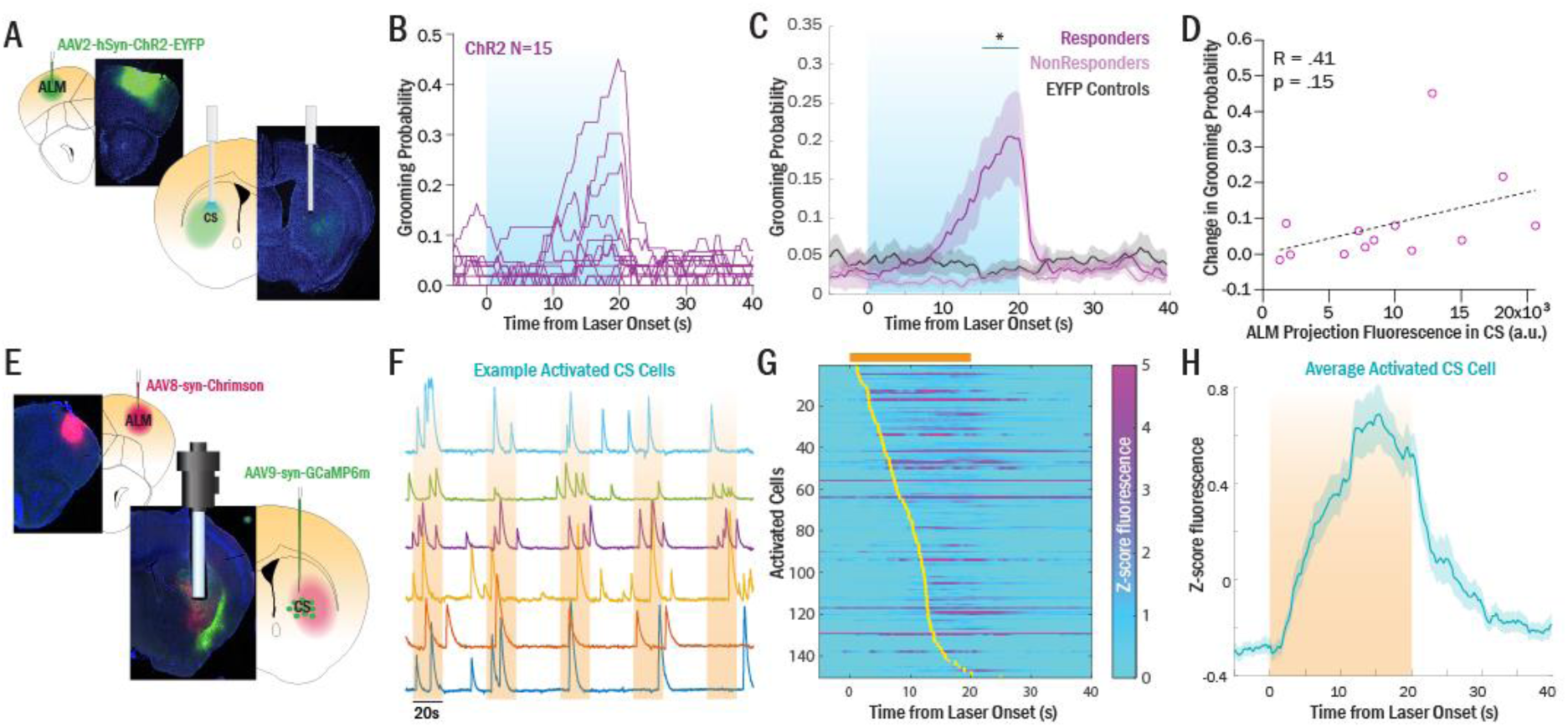
Stimulation of ALM terminals in CS causes full grooming with a latency that parallels timing of CS activation. (A) AAV5-syn-ChR2-EYFP or control AAV-5-syn-EYFP was bilaterally injected into ALM and fiberoptics were simultaneously implanted in CS. (B) Heterogeneous responses were observed following bilateral optogenetic stimulation (ChR2: N = 15). (C) Splitting the ChR2 mice into Responders (N=10) and Non-Responders (N=5) revealed a significant difference between Responders and EYFP controls (N=8, two-way repeated measures ANOVA: time x virus interaction, p < .0001, p < .05 [Sidak’s correction for multiple comparisons] for time bins 15-20s). No difference was seen between non-responders and EYFP controls (two-way repeated measures ANOVA, time x virus interaction: p = 0.009, but no significant bins with Sidak’s correction). (D) Heterogeneity in grooming response showed a non-significant correlation with heterogeneity in M2 projection intensity in CS (R^2^ = .17, p = .15. (E) nVoke 1.0 microscope (Inscopix) was used to simultaneously stimulate ALM terminals in CS (AAV8-syn-Chrimson) while recording single cell calcium activity in CS (GCaMP6m). (F) Example trace showing individual cells’ calcium response to a series of LED stimulation periods. (G) Heat map of responding cells’ average activity over 20 trials. ALM terminal stimulation evoked long latency activation responses in recorded CS cells. Yellow dots show average onset activation for each cell, calculated over 20 LED trials. (H) Average response calculated over all activated cells shows a similar temporal profile to the ALM-CS evoked grooming response (panel C).

In contrast to direct CS activation (Fig.2), ALM-CS stimulation yielded more natural-looking, complete grooming bouts (SuppVideo2). In addition, in 6/15 mice, a “stereotypy” behavior was observed, which consisted of repetitive licking of the floor or walls (SuppFig.4). However, no increases in rearing or velocity were observed (SuppFig.4). The laser-evoked grooming behavior was noticeably heterogeneous between mice (Fig.5B). To better capture this phenomenon, we therefore classified mice into grooming “responders” (N=10) or “non-responders” (N=5) based on ≥2 standard deviations increase in grooming probability over baseline mean probability during laser stimulation periods. There was a significant difference in grooming probability during laser stimulation between responders and EYFP controls (Fig.5C, repeated measures ANOVA, time x virus interaction, p < .001), but not between non-responders and EYFP controls (Fig.5C, repeated measures ANOVA, time x virus interaction p= 0.009, but no significant bins with Sidak’s multiple comparison correction). To further understand the heterogeneity in grooming response, we quantified ALM projection fluorescence in CS (Fig.5C). We observed a non-significant correlation of peak grooming probability with strength of ALM projection fluorescence in CS (R^2^ = .17, p = .15, Fig.5D), suggesting that heterogeneous activation of CS via ALM terminal stimulation may play a role in the observed heterogeneous behavioral responses.

Surprisingly, the latency to onset of behavioral responses after ALM-CS terminal stimulation was considerably delayed from stimulation onset (average latency to evoked grooming = 11.23 ± 4.24s), suggesting that ALM-evoked activation of CS may also occur at long latency. To directly investigate how ALM terminal stimulation affects CS activity, we employed unilateral simultaneous optogenetic stimulation of ALM terminals and endoscopic single-cell calcium imaging in CS (Fig.5E). Consistent with our behavioral findings, activation of ALM terminals did not immediately evoke calcium activity in most CS cells (Fig.5F-G). Instead, 78% of activated cells had an activation latency of at least 5s after LED onset (average latency in activated cells = 9.42 ± 0.39s), similar to what has recently been shown in CS during stimulation of ALM cell bodies (de Groot et al., 2020). The average activity trace of all responding cells (Fig.5H) showed temporal dynamics that matched those of the behavioral response (Fig.5B-C). These results suggest that, while artificial activation of ALM terminals is able to evoke complete grooming bouts in some circumstances, the initiation of grooming behavior may require sufficient downstream activation of CS cells.

### ALM and CS display different temporal dynamics related to grooming behavior

Together these results suggest that CS may directly initiate the movements required for grooming, while ALM activity may be required for sustaining grooming bouts. To explore this theory, we compared quantitative variables from the ALM and CS stimulation experiments. First, we compared the change in probability of grooming or grooming-like behavior as another proxy for efficacy of stimulation paradigms (Fig.6A) [activation threshold: ≥2 standard deviations over mean baseline grooming probability]. We determined that 7/9 (78%) CS-stimulation mice and 10/15 (67%) ALM-CS stimulation mice showed an evoked response (Fig.6A); these proportions were not significantly different (χ^2^ = 0.34, p = 0.56). In contrast, we observed noteworthy differences in the temporal dynamics of the evoked responses, as measured by onset latency and bout length (Fig.6B). We found that CS stimulation-evoked behavior was initiated with an average latency of 1.07s (SEM=0.07s); this was significantly earlier than the onset of the ALM-CS evoked behavioral response (12.30 ± 0.54s, t(288) = 31.93, p = 1.33×10^−96^, Fig.6C). Furthermore, bouts of grooming-like movements evoked by direct CS stimulation (4.07 ± 0.13s) were significantly shorter than more complete grooming bouts evoked by ALM terminal stimulation (6.05 ± 0.52, t(288) = 5.17, p = 4.45^-7^, Fig.6D). These data support a model in which CS activity has the capability to evoke short-latency grooming movements, but not sustain full grooming bouts. In contrast, stimulation of ALM terminals in CS evokes more qualitatively naturalistic grooming bouts that last longer, but does not immediately evoke this behavior.

**Figure 6.**
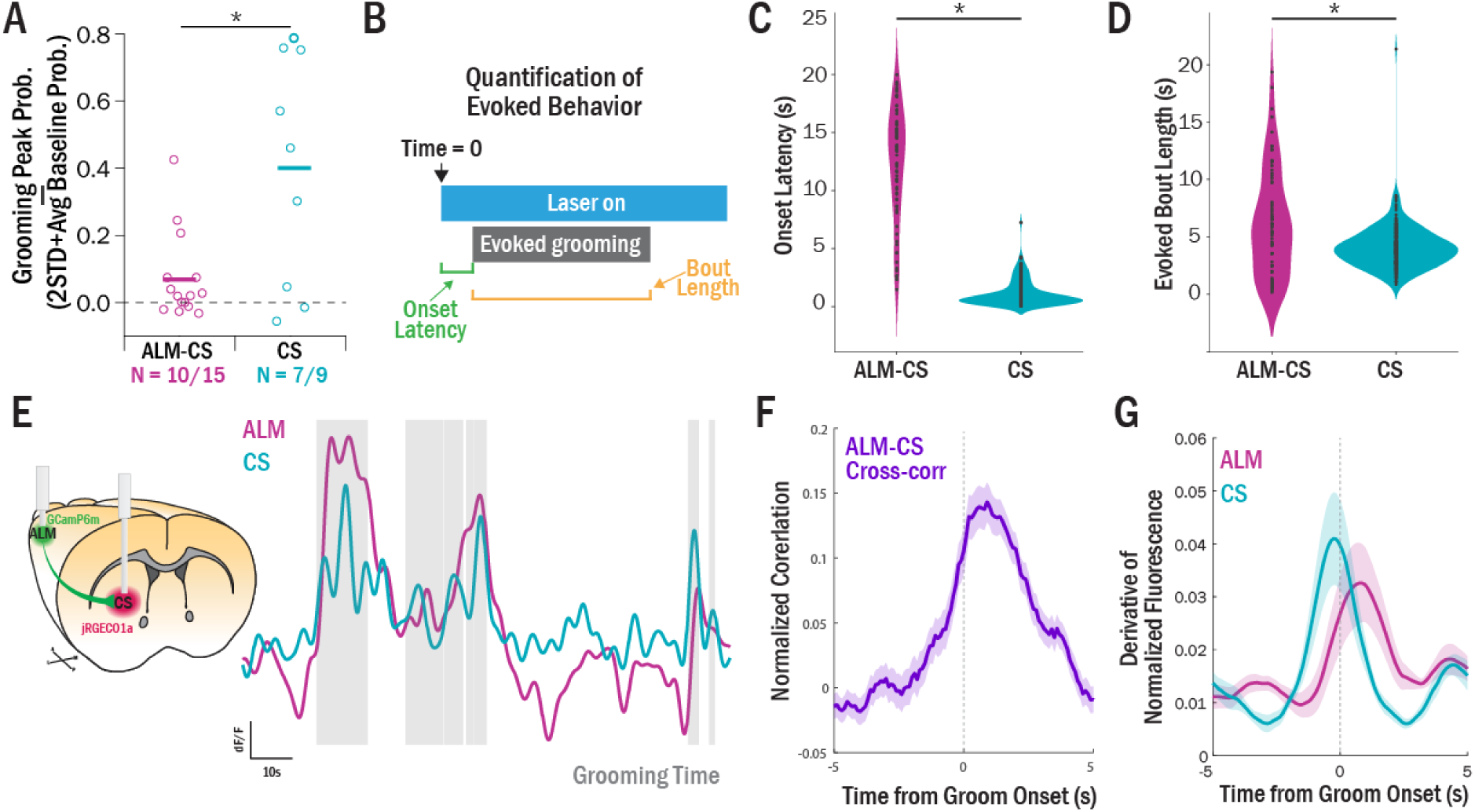
Activation of CS at grooming onset precedes ALM activation. (A) Change in grooming probability before and during laser for ALM-CS terminal stimulation and CS stimulation. As previously described, “Responders” were mice that had ≥2 standard deviations change in grooming probability from baseline. While the CS stimulation produced stronger effects overall (t-test, p = .02), the two stimulation paradigms did not have a significant difference in proportion of Responders (χ^2^ = .34, p = .56). (B) Activation dynamics were analyzed from the two stimulation paradigms. Any trials that showed either grooming fragments or full grooming movements during laser stimulation were included in this analysis. Onset latency was calculated as the time between laser onset and initiation of grooming or grooming-like behavior. Bout length was calculated as duration of an evoked grooming bout that occurred during laser-on time. (C) Onset latency of grooming was significantly earlier in CS stimulation trials relative to ALM-CS terminal stimulation trials (t(288) = 31.93, p = 1.33×10^−96^). (D) Duration of evoked grooming bouts was significantly longer in ALM-CS terminal stimulation trials relative to CS stimulation trials (t(2 88) = 5.17, p = 4.45^-7^). (E) Dual-color photometry was conducted using GCaMP6m in ALM and jRGECO1a in CS, with fiber optics implanted unilaterally above the virus injection on the same side. Example traces show similarly-varying signals in ALM and CS. (F) Cross-correlation of the simultaneously-recorded signals at grooming onset showed that CS precedes ALM (comparison of area under the curve on either side of time=0, t(174) = -4.61, p = 7.63×10^−6^). (G) Derivatives of each signal were calculated to better compare across different indicator kinetics. Aligning the derivative signal in ALM or CS to grooming onset showed that the greatest rate of change in CS occurs before grooming onset, while the greatest change in ALM activity occurs after grooming onset (significant difference in peak time between ALM and CS, WRST, p=0.004).

To further test this model, we simultaneously recorded calcium activity in ALM (GCaMP6m) and CS (jRGECO1a) during spontaneous grooming behavior (Fig.6E). Visual observation of the two time-varying signals showed similar changes in activity across the two regions (Fig.6E). To characterize the relationship between activity in these two regions, we calculated the cross-correlation between the ALM and CS calcium signals at grooming onset (Fig.6F). Quantification of the area under the curve for each side showed that the cross-correlation was significantly weighted towards the right (t(174) = -4.61, p = 7.63×10^−6^), indicating that CS activity leads ALM activity at grooming onset.

This indication that CS precedes ALM in grooming-relevant activity was surprising given that cortical inputs are thought to drive striatal activity. Thus, we conducted an additional test on the simultaneous signals to confirm our results. First, we performed a derivative transformation which has been shown to be an estimation of spiking activity (Markowitz et al., 2018) to provide a direct comparison of the rates of change of each signal. This transformation was then aligned to grooming onset. Paralleling our initial findings, we observed that CS exhibited the highest rate of change immediately before grooming, i.e., shows its primary activity increase immediately prior to grooming onset (Fig.6G). In contrast, ALM displayed the highest rate of change following grooming onset, consistent with our initial findings that ALM activity starts to increase after groom initiation (Fig.6G). Among simultaneously recorded animals, there was a significant difference between the time of peak change in ALM (1.46 ± 0.59s *after* grooming onset) and CS activity (1.99 ± 1.30s *before* grooming onset) (WRST p=0.004). These experiments provide additional evidence that CS activity precedes ALM activity in the initiation of grooming behavior.

## Discussion

Our data reveal that the ALM-CS circuit plays a prominent role in evoking and sustaining spontaneous grooming behavior in mice. First, we found that increases in CS activity are associated with the start of grooming bouts, and not other spontaneous behaviors. In addition, mimicking this activity in CS is sufficient to evoke short, fragmented movements that resemble grooming behavior. Next, we found that anterior lateral motor area (ALM), a prominent input to CS, also shows increased activity at groom onset; however, this activity ramps up during each bout, with a peak that is highly correlated with grooming bout length. Furthermore, optogenetically activating ALM projections in CS evokes full grooming behavior with a long latency that parallels the time course of CS activation. Finally, analysis of simultaneous recordings in ALM and CS demonstrates that the increase in CS activity leads the increase in ALM activity at grooming onset. Taken together, these data indicate that striatal activation precedes cortical activation to initiate naturalistic grooming behavior, and suggest a general model in which striatum initiates spontaneous movements, while reverberating activity in cortico-basal ganglia-thalamic loops is required to properly sequence and sustain them.

### Specialization of ALM-CS circuit for grooming may partially reflect its role in licking behavior

Our results indicate that the ALM-CS circuit is relatively specific for grooming behavior, and appears to be particularly associated with body grooming. We demonstrated that CS and ALM had greater increases in calcium activity associated with body grooming than with face grooming (Fig.1, Fig.3), and our optogenetic stimulation paradigms often produced behaviors that resembled body grooming (turning inward towards the body and performing tongue movements), as opposed to face grooming (raising paws to face). This selectivity of CS for body grooming may reflect the known role of ALM in anticipatory and ingestive licking behavior in trained tasks that require consumption of liquid (Bollu et al., 2019; Li et al., 2015). Here, in an untrained context, we observed grooming and grooming-like behaviors that also involved licking. This convergent data suggests that, in mice, the ALM-CS circuit may have evolved to be selective for licking behavior, which can be applied in several different contexts critical for mouse survival, including behaviors critical for hygiene and consumption of food and fluid.

### CS may uniquely integrate motor, cognitive, and limbic information to generate species-typical behaviors essential for survival

Though it is somewhat surprising that activation of the CS results in a very specific type of movement, these findings are reminiscent of classic topographical models of the striatum generated based on data from the primate literature. For instance, striatal disinhibition via bicuculline causes very different behavioral effects depending on which subregion is injected (Worbe et al., 2008). While disinhibition of dorsolateral striatum caused hyperactivity and dyskinetic movements of specific body parts, bicuculline injections into more central regions of striatum caused stereotypies, including perseverative grooming (Worbe et al., 2008). Thus, our data are broadly consistent with the idea that the striatum plays a key role in the generation of movements, and that the particular movement generated may be dictated by striatal topography.

As discussed above, the ability of the CS to generate grooming movements suggests that this region may be specialized for behaviors critical for survival. Interestingly, in rodents, CS receives input from several key regions in addition to ALM, including fore- and hind-limb motor cortical regions (Ebrahimi et al., 1992) and associative orbitofrontal cortex (Burguiere et al., 2013; Corbit et al., 2019). Therefore, CS is able to integrate information about several different body parts, as well as information about action value that can be used to adapt behavioral responses. This unique convergence of inputs could explain why CS may be important for self-regulating ethologically-important behaviors, like grooming, that are highly related to an animal’s level of stress and anxiety in a particular environment (Fernández-Teruel and Estanislau, 2016).

### CS activation generates movement fragments

Our data show that activity in CS is sufficient to produce short-latency grooming-like movements, and therefore suggest that activity in CS may directly drive the initiation of naturalistic grooming behavior via the generation of movement fragments. Specifically, though our photometry data indicate that CS activity is associated with initiation of grooming behavior in naturalistic contexts, artificial optogenetic CS stimulation produces fragmented and repetitive components of grooming, rather than smoothly sequenced grooming bouts. Importantly, these data support the broad possibility that specific movement syllables are represented by unique activity patterns in CS, consistent with past work showing that disinhibition of CS causes stereotyped tic-like behavior (Pogorelov et al., 2015). This is in contrast to prior work in DLS and DMS that has shown similar activation profiles for several different types of movements, including grooming (Barbera et al., 2016; Cui et al., 2013; Markowitz et al., 2018; Parker et al., 2018). This divergence in results suggests that CS may represent more specific behavioral repertoires crucial for survival that are linked to central pattern generators, whereas DLS/DMS may instead represent activation of specific muscles and postures that are widely used across multiple behaviors.

### ALM may facilitate smooth, sustained movements through appropriate sequencing of behavior syllables

How do movement fragments become organized into smoothly sequenced behaviors? Our data support a corticostriatal model of action selection in which ALM contributes to the organization and sustainment of sequences of behavioral syllables that are initiated by CS. A role for ALM in generating behavioral sequences is supported by previous work showing that supplementary motor regions contribute to trained sequenced behavior (Gaymard et al., 1990; Mushiake et al., 1990; Rothwell et al., 2015). Consistent with a role for ALM in linking together a series of movement syllables, our data suggest that sustained ramping activity in ALM encodes the length of a self-initiated grooming bout. Similar ramping activity has been shown in ALM between a sample and the subsequent lick response in a delay-match-to-sample task (Guo et al., 2014; Inagaki et al., 2018). Furthermore, supplementary motor regions in humans have been shown to have activity related to the duration of a waiting period in a timed motor production task (Macar et al., 2004; Macar et al., 2006; Macar et al., 1999).

However, our results are in contrast to previous findings that supplementary motor areas show preparatory activity before a movement (Deecke, 1987; Guo et al., 2014; Li et al., 2015; Mita et al., 2009; Roland et al., 1980). One important distinction between our results and this prior work is that previous studies all looked at trained tasks in which the anticipated response was well-known. Thus, the temporal coding observed in these studies was associated with a sustained waiting period between a cue and a movement (Deecke, 1987; Li et al., 2015; Macar et al., 2004; Macar et al., 2006; Macar et al., 1999; Mita et al., 2009). This is likely to be a fundamentally different process from the generation of spontaneous, internally-generated movements. One theory that could merge these seemingly disparate findings is that supplementary motor areas encode the sustainment of various behavioral states, including continuous repetitive movement (naturalistic grooming) or prolonged waiting postures (periods between trained cues and responses). Similar to our findings that longer grooming bouts showed greater ALM activation, work in primates supports this theory by showing that longer waiting periods evoke greater activity in pre-supplementary motor area (Mita et al., 2009). Taken together, these data suggest that supplementary motor regions across species are essential for temporal encoding of sustained behavioral sequences or waiting periods.

### A model of grooming generation in the ALM-CS circuit

Taken together, our data support a model in which grooming syllables are initiated through striatal activation, but sequencing and sustainment of these syllables into normal grooming motifs is encoded by ALM. Specifically, we show that increases in CS activity precede increases in ALM activity during the initiation of spontaneous grooming bouts. Furthermore, optogenetic and photometry experiments demonstrate that CS activity is more directly related to the initiation of grooming behavior than ALM. This suggests that CS activation is a driver of grooming initiation, and that ALM may be activated after grooming initiation to support sequencing of movement fragments. This is in contrast to classic models that posit that action plans are represented in cortex and gated through striatum (Frank, 2011). A potential circuit to support this mechanism exists: the downstream output nucleus of the striatum, the substantia nigra, projects to ventromedial (VM) thalamus (Deniau and Chevalier, 1985), a region which has recently been shown to have a direct effect on ALM activity (Guo et al., 2017). Specifically, activity in this thalamic region has been shown to support persistent activity in ALM (Guo et al., 2017). Thus, one possibility for how the ALM-CS circuit may generate continuous grooming bouts is via recurrent activity in this ALM-CS-VM-thalamic loop. Specifically, CS activity could initiate grooming, and downstream VM thalamus activation could then cause reverberant, persistent activity in ALM to sustain grooming.

A remaining question unanswered by our data is which region(s) are responsible for terminating grooming via inhibition of ALM ramping activity. One candidate region is the subthalamic nucleus (STN), which has been associated with stopping movements (Adam et al., 2020; Aron and Poldrack, 2006; Schmidt et al., 2013). As it is known to receive direct inputs from cortex (Afsharpour, 1985) and thought to have a downstream effect of inhibiting thalamo-cortical transmission (Aron and Poldrack, 2006), the STN is well-placed to potentially serve as a terminator of grooming behavior. For example, the ALM activity peak towards the end of a grooming bout could cause STN to cross an activation threshold, leading to dampening of thalamo-cortical transmission and termination of grooming. Supporting this hypothesis, it has recently been shown that activation of posterior ALM projections to STN is sufficient to cause termination of ongoing locomotion (Adam et al., 2020). However, this work investigated externally-triggered stops, rather than internally generated terminations of behavior. To investigate the possibility that STN may play a role in terminating spontaneous movements as well, substantial work must be done to investigate the role of STN in naturalistically-generated movements.

### Conclusions

These data present a corticostriatal framework for understanding how spontaneous behavioral components may be initiated and sequenced into smooth movements. Further, our results suggest that supplementary motor regions may be a useful target for treatment of abnormal sequencing and sustainment of behaviors in illnesses such as Obsessive-Compulsive Disorder (OCD). Consistent with this idea, pre-supplementary motor area (pre-SMA) has been identified as a promising target for transcranial magnetic stimulation treatment in OCD (Berlim et al., 2013; Mantovani et al., 2010). In contrast, disorders characterized by generation of abnormal movement fragments may be better treated by targeting striatum and downstream regions such as thalamus or STN, which have been identified as promising targets for Tourette Syndrome and dyskinesia (Viswanathan et al., 2012; Welter et al., 2010).

## Methods

### Animals

Male and female wild-type (WT) C57BL6 background mice were used for all experiments. Most WT mice were genotype-confirmed littermates of Sapap3-KO mice, a colony initially established at MIT by Dr. Guoping Feng. One cohort of C57BL6/J WT mice (part of ALM-CS Stim, Fig.5) was purchased directly from Jackson Laboratory. Mice were group housed with 2-5 mice per cage except when noted. All mice had *ad libitum* access to food and water. Animals were randomly assigned to experimental or control groups, balancing for sex and cage mates. All experiments were approved by the Institutional Animal Use and Care Committee at the University of Pittsburgh in compliance with National Institutes of Health guidelines for the care and use of laboratory animals.

### Stereotaxic Surgeries

Mice underwent stereotaxic surgery between the ages of 4 and 8 months. Stereotaxic surgeries were performed under isofluorane anesthesia (2%). Burr holes were drilled over the target location for subsequent virus injection or implant. Virus was injected using a syringe pump (Harvard Apparatus) fitted with a syringe (Hamilton) connected to PE10 tubing and a 30 gauge cannula and allowed to incubate for at least 3 weeks before experiments.

All recording (single-cell calcium imaging, fiber photometry) experiments were conducted using unilateral virus injections and implants. For fiber photometry, AAV9-Synapsin-GCaMP6m-WPRE-SV40 (250nL, Addgene) was injected into ALM (AP 2.90, ML 1.55, DV .75mm) and/or AAV1-syn-NES-jRGECO1a-WPRE-SV40 (500nL, Addgene) was injected into CS (AP .50, ML 1.95, DV 3.00mm). Optical fibers (NA = .37) were implanted into ALM and CS at the same AP and ML coordinates, but were 0.15mm above the injection site. For combined optogenetic stimulation and calcium imaging experiments, AAV8-syn-ChrimsonR-tdT (300nl, Addgene) was injected into ALM and AAV9-hSyn-GCaMP6m (800nl, Addgene) was injected into CS. A 6mm long 500um diameter GRIN lens was implanted over the injection site in CS, to simultaneously stimulate ALM terminals and record calcium activity from CS soma.

All optogenetic behavioral manipulations were conducted with bilateral virus injections of AAV2-hSyn-ChR2-EYFP (500nL, Addgene) into either ALM (AP 2.90, ML 1.55, DV 0.75mm) or CS (AP 0.70, ML 2.00, DV 3.00mm, fibers at 2.60-2.85mm). For ALM terminal stimulation, AAV2-hSyn-ChR2-EYFP (350nL, Addgene) was injected into ALM and fibers were implanted into CS (AP 0.70, ML 2.00, DV 2.60-2.85mm). For CS stimulation, fibers were implanted 0.15-0.40mm above the viral injection site (AP 0.70, ML 2.00, DV 2.60-2.85mm).

### Optogenetic Behavioral Manipulations

After 4-6 weeks of virus incubation and recovery, mice were handled for several days prior to behavior experiments. All mice were habituated to the observation chamber and optical fiber tethering for three days prior to behavioral manipulation. On experiment day, mice were scruffed and attached to optical cables and placed in a 10×10 inch clear plexiglass observation chamber. A Point Grey camera was fixed beneath the chamber and behavior was filmed from below. For CS stimulation experiments, 5mW 470nm light was used. Fifty-one 10s trials of constant light were presented with a pseudorandom inter-trial interval with an average of 30s (25-35s, 5s jitter). All experimenters were blinded to experimental condition.

Stimulation of ALM terminals was altered due to the propensity of cortical optogenetic stimulation to cause seizure activity. 20Hz (10ms pulse width) pulsed 470nm light was used. Initial experiments were conducted at 10mW light power, and animals were monitored for seizure or pre-seizure activity. For animals that did show seizure activity, light was lowered first to 7mW and then to 5mW if seizure activity persisted. Sessions included in analysis were the highest light power used that did not cause seizure activity; additionally, any trials within the session that had pre-seizure activity were excluded from analysis. Pilot testing showed that 1) 20s light periods were more likely to show grooming behavior than 10s light periods, and 2) grooming behavior was reduced as trial number increased (SuppFig.3). Thus, data are presented from experiments conducted with 20s light pulses, and only the first usable 20 trials (e.g. without pre-seizures) of this experiment were analyzed.

Behavioral data were manually scored by blinded raters using Noldus Observer. Grooming probability was binned into 500ms time bins and analyzed with two-way repeated-measures ANOVAs and post-hoc t-tests using Sidak’s p-value correction (Prism, Graphpad).

### Fiber Photometry

Fiber photometry experiments were conducted in freely behaving mice in a 10×10 inch clear plexiglass chamber. A Neurophotometrics 3-color, 2-site system was used to collect imaging data (Neurophotometrics). Three LEDs (415nm, 470nm, 560nm) were pulsed at 30Hz in an interleaved manner to obtain 1) isosbestic motion signal, 2) GCaMP6m activity, and 3) jRGECO1a activity. The recorded trace was then separated to obtain activity for each channel individually.

After separating the 3 channels, a linear fit of the isosbestic signal to the activity channels (470nm or 560nm) was calculated. This linear fit was then subtracted from each corresponding activity channel to remove baseline fluorescence and motion artifacts. An additional moving minimum baseline (2min sliding window) was subtracted from each resulting trace to account for slow fluctuations in activity, such as additional decay from bleaching. Finally, each activity trace was normalized by dividing by the standard deviation. Manually (grooming, rearing) or automatically (locomotion) scored behaviors were then aligned to the activity traces using an initial “session start” LED signal for alignment.

Unless otherwise noted, analyzed grooming bouts were restricted to bouts that did not show grooming for 3s prior to grooming bout onset. Trials were then zeroed to the 3s baseline period by subtracting the mean activity in that interval from the overall trace. Statistical analysis of time-varying calcium traces was performed using two-way repeated-measures ANOVAs with post-hoc contrasts comparing the activity at 1s before behavior onset to all other time bins (SPSS, IBM). All statistical tests were conducted across individual animals’ average trial data unless otherwise noted.

For peak detection analysis, grooming trials were further excluded if they had any additional grooming initiations in the 10s following grooming bout onset, to avoid grooming onset contamination in the detection of peaks. Peak analysis was conducted using the MATLAB function *findpeaks*. This function finds the local maxima in a trace and provides information about time, amplitude, and “prominence”– a measure that takes into account the amplitude of a given peak, the slopes on either side, and the amplitude of the peaks surrounding the peak of interest. The first peak reaching the prominence threshold that occurred after grooming onset was the automatically detected peak. Time was extracted relative to grooming onset time for each trial, and absolute amplitude was calculated from the baselined trials.

Four cohorts of mice were used in the collection of photometry data (Cohort 1: GCaMP6m in ALM, total N=9 with ALM signal; Cohort 2: GCaMP6m in ALM, JRGECO1a in CS, total N = 3, 1 with CS signal; Cohort 3: GCaMP6m in ALM, JRGECO1a in CS, total N = 7, 3 with CS signal; Cohort 4: GCaMP6m in ALM, JRGECO1a in CS, total N = 5, 5 with CS signal). Reasons for no signal in some animals include poor targeting of virus and/or fiber, and fibers being pulled out of the cement headcap. All mice that had CS signal were used in Figure 1 (N=9). The ALM-only imaging cohort (Cohort 1) was used for data in Figure 3. To gain additional data for the bout length analyses (Figure 4), additional ALM-GCaMP6m trials from the dual-color photometry cohorts were added to the analysis (Cohorts 2-4). Finally, for the dual-color photometry analyses (Figure 6), any animal with signal in both ALM and CS was used (N = 6/15).

### Freely moving microendoscopy

Mice were habituated to microscope attachment and the grooming chamber for three consecutive days prior to testing. For recording single cell activity in the CS, analog gain of the image sensor was set between 1 and 4 while the 470 nm LED power was set between 10 to 30% transmission range. Stimulation of ALM terminals infected with ChRimson was achieved through the delivery of 600nm amber light through the objective lens as GCaMP-positive cells are simultaneously being excited with 460nm blue light (Stamatakis et al., 2018). After mice were placed into the chamber and calcium imaging began, the 600 nm optogenetic LED (OG-LED) was turned on. OG-LED stimulation consisted of 15 pulse trains of 20 Hz stimulation for 20s. Each 20 Hz train was followed by a 30s interval of no stimulation. Each session lasted 15 minutes in total.

Following acquisition, raw calcium videos were spatially downsampled by a binning factor of 4 (16x spatial downsample) and temporally downsampled by a binning factor of 2 (down to 10 frames per second) using Inscopix Data Processing Software (v1.3.0, Inscopix Inc, Palo Alto, CA USA). Lateral brain motion was corrected using the registration engine TurboReg (Ghosh et al., 2011), which uses a single reference frame to match the XY positions of each frame throughout the video. Motion corrected 10 Hz video of raw calcium activity was then saved as a .TIFF and used for cell segmentation.

Using custom MATLAB scripts, the motion corrected .TIFF video was then processed using the Constrained Non-negative Matrix Factorization approach (CNMFe), which has been optimized to isolate signals from individual putative neurons from microendoscopic imaging (Zhou et al., 2018). The CNMFe method is able to simultaneously denoise, deconvolve, and demix imaging data (Pnevmatikakis et al., 2016) and represents an improvement over previously used algorithms based on principle component analysis (Zhou et al., 2018). Putative neurons were identified and manually sorted according to previously established criteria (Pnevmatikakis et al., 2016). For each individual cell, the raw fluorescence trace was Z-scored to the average fluorescence and standard deviation of that same trace. Thus, fluorescence units presented here are referred to as “Z-scored fluorescence”.

To identify central striatal cells modulated by ALM terminal activation, the Z-scored fluorescence response for each cell to 20s of OG-LED stimulation was averaged across presentations (15 total presentations). This average was then compared with an unpaired *t*-test to the periods immediately preceding (10s before) and during (20s) stimulation (Bonferroni multiple comparison correction *p* ≤ 0.0001).

### Histology

After experiments were completed, mice were transcardially perfused using 4% paraformaldyhyde (PFA) and post-fixed in PFA for 24 hours. Post-hoc confirmation of viral and implant targeting was conducted on 35um slices from the harvested brains. Slices were mounted with DAPI coverslipping media and inspected for relevant fluorophores (e.g. GFP or mCherry). Fiber implants were identified by finding damage tracks in the brain at the specific location.

## Supplementary Figures

**Supplementary Figure 1.**
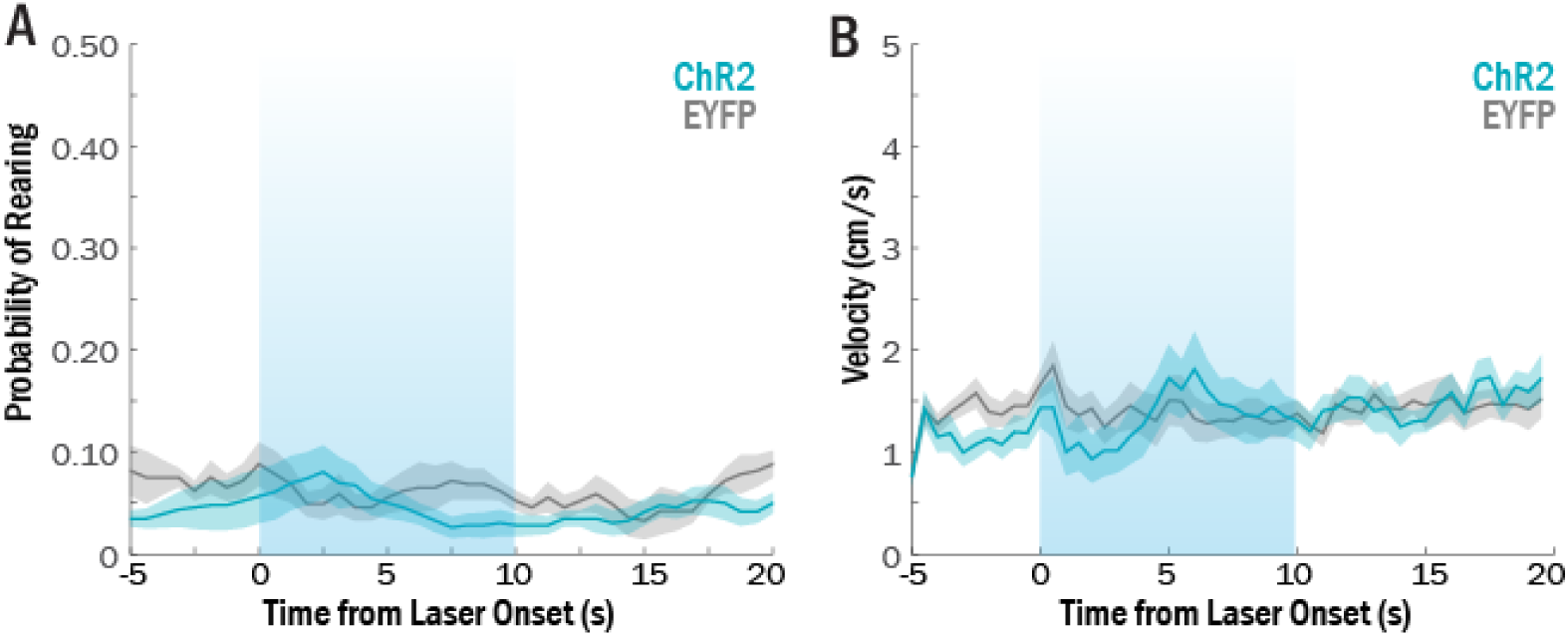
Optogenetic stimulation of CS does not cause increased rearing or change in velocity. (A) Bilateral 10s stimulation of CS did not increase probability of rearing behavior (ChR2: N = 9; EYFP: N = 6, two-way repeated-measures ANOVA, time x virus interaction: p = .26). (B) Stimulation of CS did not cause changes in velocity (two-way repeated-measures ANOVA, time x virus interaction: p = .88).

**Supplementary Figure 2.**
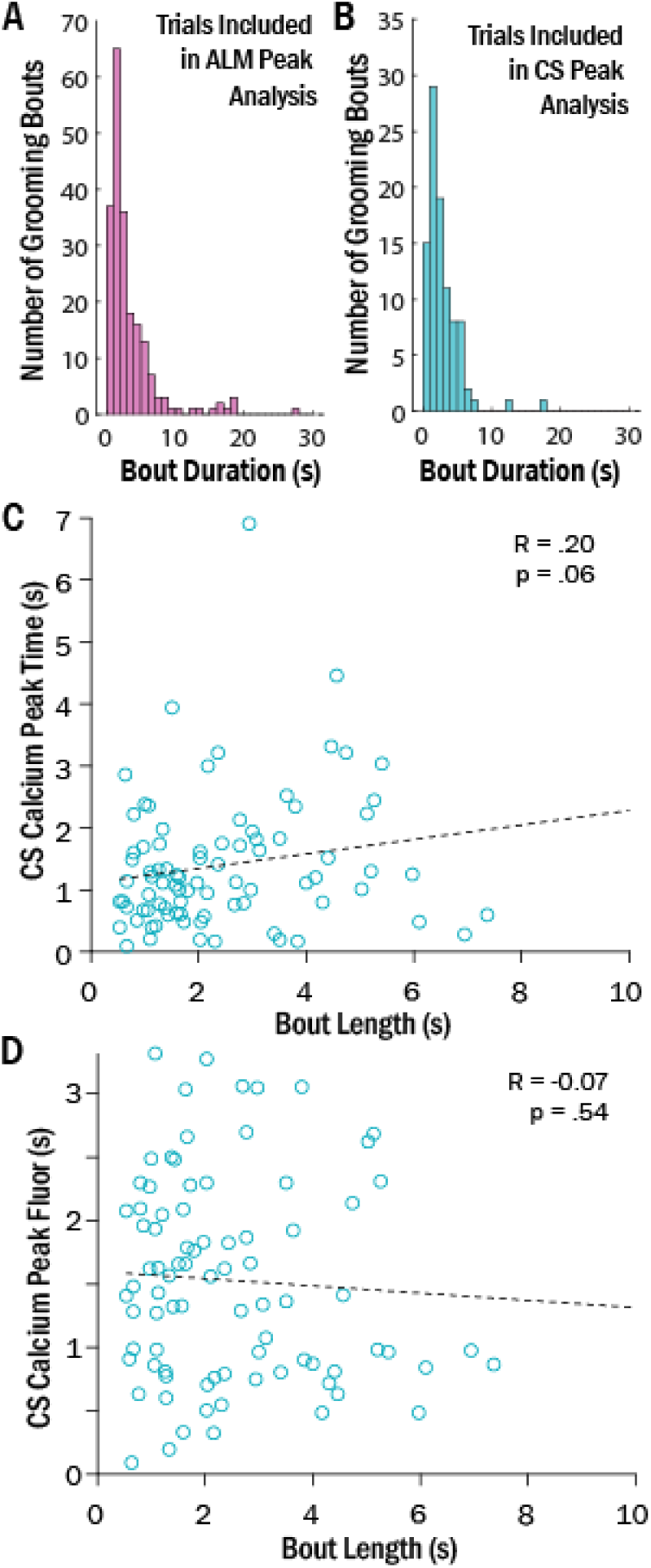
CS peaks do not correlate with bout length. (A) Histogram showing distribution of bout durations of the trials included in ALM peak analysis (Fig.4). (B) Histogram showing distribution of bout durations of trials included in CS peak analysis. Distributions are not significantly different (Kolmogorov-Smirnov test, p = .87). (C) CS activity peak time plotted against the bout duration for a given trial. A non-significant, trend correlation exists between peak time and bout length (R^2^ = .40, p = .06). (D) CS activity amplitude of detected peak plotted against bout duration for a given trial. No significant correlation was detected (R^2^ = .005, p = .54).

**Supplementary Figure 3.**
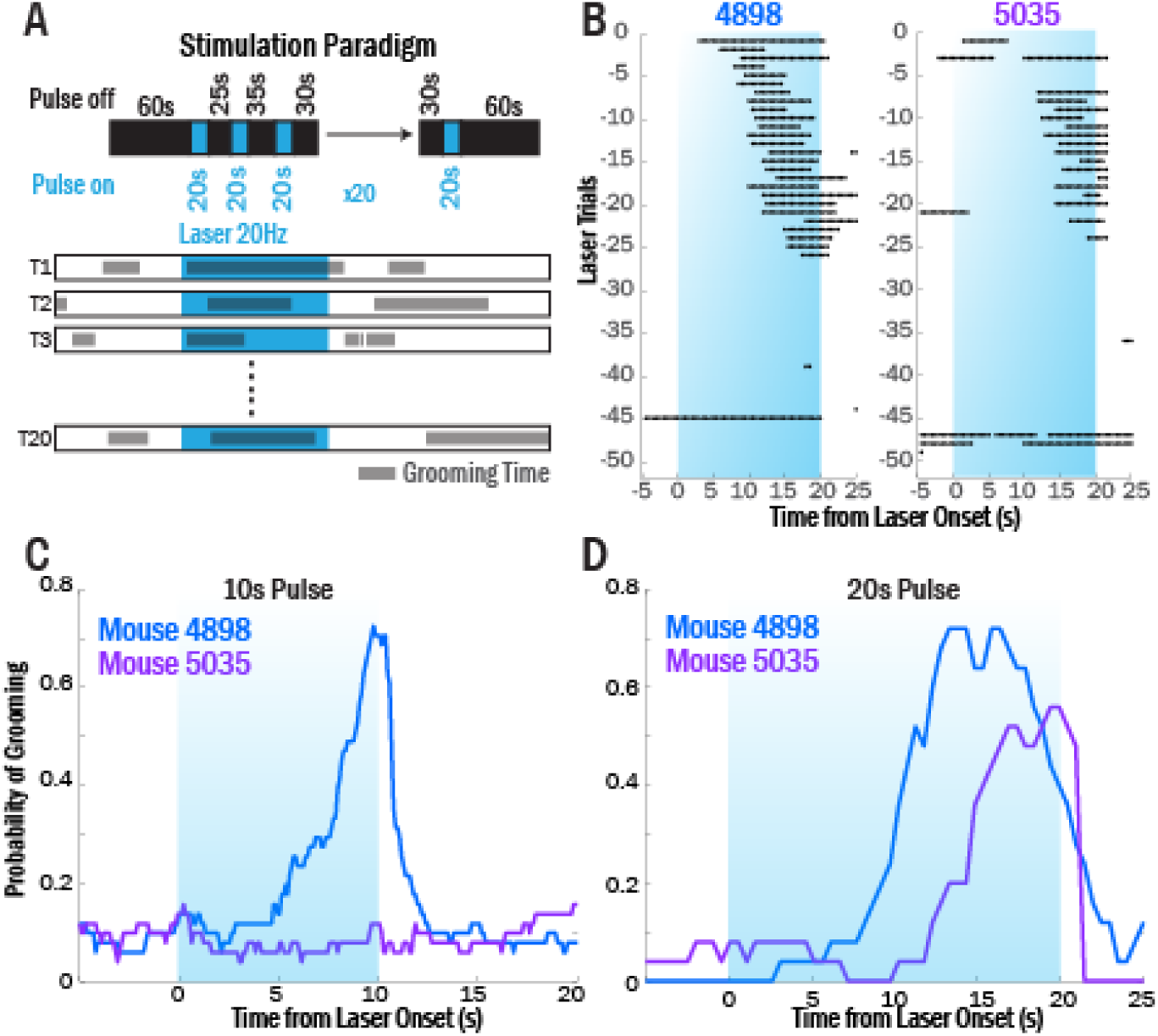
Pilot ALM-CS stimulation experiments show longer-latency responses relative to CS stimulation, and effects that deteriorated over the session. (A) Schematic showing the stimulation paradigm used for data displayed in Figure 5. Laser was pulsed at 20Hz (10ms pulses) for 20s periods, with an inter-trial interval of 25-35s. The first 20 trials were used in the calculation of behavior probability. (B) Two example mice from a pilot cohort (ChR2: N = 5; EYFP: N = 3) that showed decreased grooming response to stimulation as the experiment progressed, with no grooming observed after trial ∼20-25. Based on these findings, only the first 20 trials were analyzed. (C) Two example mice from the first pilot experiment with 10s pulses, showing that only mouse 4898 showed a grooming response with this laser duration. (D) When laser periods were lengthened to 20s, the second mouse (5035) also showed a grooming response that started after 10s.

**Supplementary Figure 4.**
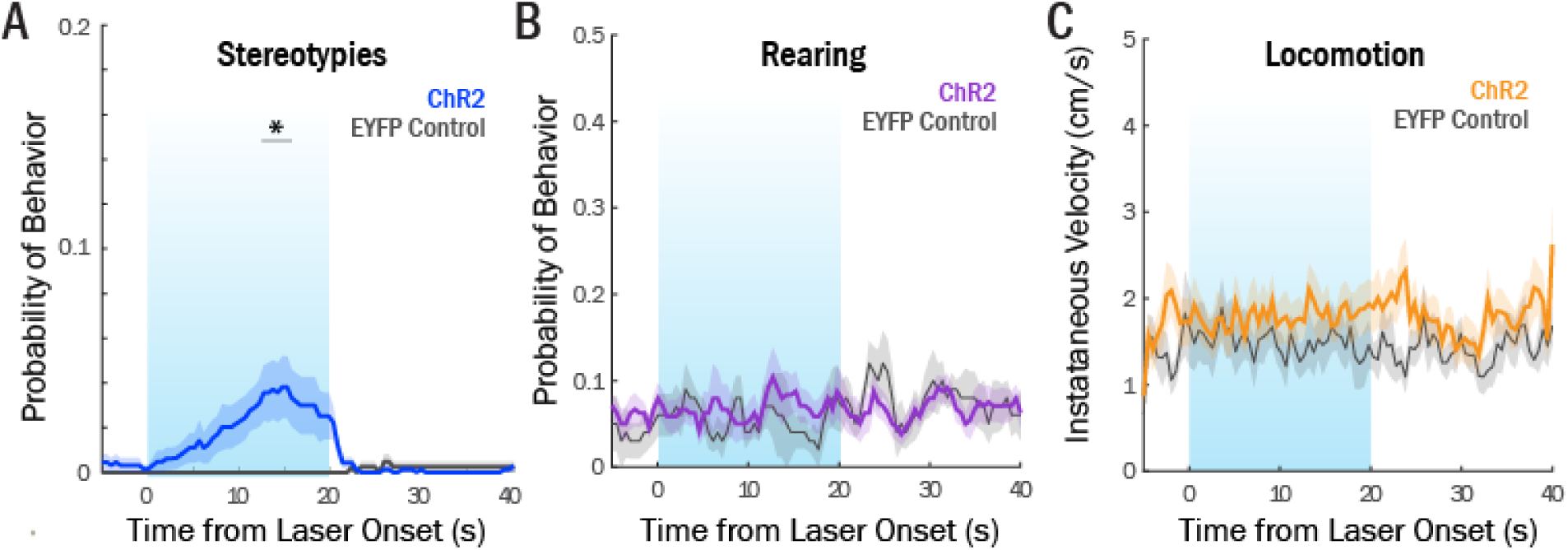
ALM-CS stimulation causes a heterogeneous response of grooming and stereotypies, but does not cause rearing or locomotion. (A) Group averages of peri-laser stereotypy probability for ChR2 (N=15) and EYFP control mice (N=8, two-way repeated measures ANOVA, time x virus interaction: p < .0001, bins 12.5-15.5s significant with Sidak’s multiple comparison correction). (B) Group averages of peri-laser rearing probability for ChR2 (N=13) and EYFP control mice (N=5, two-way repeated measures ANOVA, time x virus interaction: p = .96). (C) Group averages of peri-laser velocity for ChR2 (N=13) and EYFP control mice (N=5, two-way repeated measures ANOVA, time x virus interaction: p = .17).

## References

1. Adam, E.M., Johns, T., and Sur, M. (2020). Cortico-subthalamic projections send brief stop signals to halt visually-guided locomotion. bioRxiv.

2. Afsharpour, S. (1985). Topographical projections of the cerebral cortex to the subthalamic nucleus. Journal of Comparative Neurology 236, 14–28.

3. Alexander, G.E., DeLong, M.R., and Strick, P.L. (1986). Parallel organization of functionally segregated circuits linking basal ganglia and cortex. Annu Rev Neurosci 9, 357–381.

4. Aron, A.R., and Poldrack, R. (2006). Cortical and subcortical contributions to stop signal response inhibition: role of the subthalamic nucleus. Journal of Neuroscience 26, 2424–2433.

5. Balleine, B.W., and O’Doherty, J.P. (2010). Human and rodent homologies in action control: corticostriatal determinants of goal-directed and habitual action. Neuropsychopharmacology 35, 48.

6. Barbera, G., Liang, B., Zhang, L., Gerfen, C.R., Culurciello, E., Chen, R., Li, Y., and Lin, D.-T. (2016). Spatially compact neural clusters in the dorsal striatum encode locomotion relevant information. Neuron 92, 202–213.

7. Berlim, M.T., Neufeld, N.H., and Van den Eynde, F. (2013). Repetitive transcranial magnetic stimulation (rTMS) for obsessive–compulsive disorder (OCD): An exploratory meta-analysis of randomized and sham-controlled trials. Journal of psychiatric research 47, 999–1006.

8. Bollu, T., Whitehead, S.C., Kardon, B., Redd, J., Liu, M.H., and Goldberg, J.H. (2019). Tongue Kinematics. Cortex-dependent corrections as the mouse tongue reaches for, and misses, targets. bioRxiv, 655852.

9. Burguiere, E., Monteiro, P., Feng, G., and Graybiel, A.M. (2013). Optogenetic stimulation of lateral orbitofronto-striatal pathway suppresses compulsive behaviors. Science 340, 1243–1246.

10. Corbit, V.L., Manning, E.E., Gittis, A.H., and Ahmari, S.E. (2019). Strengthened inputs from secondary motor cortex to striatum in a mouse model of compulsive behavior. Journal of Neuroscience 39, 2965–2975.

11. Cui, G., Jun, S.B., Jin, X., Pham, M.D., Vogel, S.S., Lovinger, D.M., and Costa, R.M. (2013). Concurrent activation of striatal direct and indirect pathways during action initiation. Nature 494, 238.

12. de Groot, A., van den Boom, B.J., van Genderen, R.M., Coppens, J., van Veldhuijzen, J., Bos, J., Hoedemaker, H., Negrello, M., Willuhn, I., and De Zeeuw, C.I. (2020). NINscope, a versatile miniscope for multi-region circuit investigations. eLife 9, e49987.

13. Deecke, L. (1987). Bereitschaftspotential as an indicator of movement preparation in supplementary motor area and motor cortex. Paper presented at: Ciba Found Symp (Wiley Online Library).

14. Deniau, J., and Chevalier, G. (1985). Disinhibition as a basic process in the expression of striatal functions. II. The striato-nigral influence on thalamocortical cells of the ventromedial thalamic nucleus. Brain Research 334, 227–233.

15. Ebrahimi, A., Pochet, R., and Roger, M. (1992). Topographical organization of the projections from physiologically identified areas of the motor cortex to the striatum in the rat. Neuroscience research 14, 39–60.

16. Fernández-Teruel, A., and Estanislau, C. (2016). Meanings of self-grooming depend on an inverted U-shaped function with aversiveness. Nature Reviews Neuroscience 17, 591.

17. Frank, M.J. (2011). Computational models of motivated action selection in corticostriatal circuits. Current opinon in neurobiology 21, 381–386.

18. Gaymard, B., Pierrot-Deseilligny, C., and Rivaud, S. (1990). Impairment of sequences of memory-guided saccades after supplementary motor area lesions. Annals of Neurology: Official Journal of the American Neurological Association the Child Neurology Society 28, 622–626.

19. Ghosh, K.K., Burns, L.D., Cocker, E.D., Nimmerjahn, A., Ziv, Y., El Gamal, A., and Schnitzer, M.J. (2011). Miniaturized integration of a fluorescence microscope. Nature Methods 8, 871.

20. Graybiel, A.M. (1998). The basal ganglia and chunking of action repertoires. Neurobiology of Learning and Memory 70, 119–136.

21. Guo, Z.V., Inagaki, H.K., Daie, K., Druckmann, S., Gerfen, C.R., and Svoboda, K. (2017). Maintenance of persistent activity in a frontal thalamocortical loop. Nature 545, 181.

22. Guo, Z.V., Li, N., Huber, D., Ophir, E., Gutnisky, D., Ting, J.T., Feng, G., and Svoboda, K. (2014). Flow of cortical activity underlying a tactile decision in mice. Neuron 81, 179–194.

23. Haber, S.N. (2016). Corticostriatal circuitry. Neuroscience in the 21st Century, 1–21.

24. Inagaki, H.K., Inagaki, M., Romani, S., and Svoboda, K. (2018). Low-dimensional and monotonic preparatory activity in mouse anterior lateral motor cortex. Journal of Neuroscience 38, 4163–4185.

25. Jin, X., Tecuapetla, F., and Costa, R.M. (2014). Basal ganglia subcircuits distinctively encode the parsing and concatenation of action sequences. Nature neuroscience 17, 423.

26. Kim, J., and Ragozzino, M.E. (2005). The involvement of the orbitofrontal cortex in learning under changing task contingencies. Neurobiology of Learning and Memory 83, 125–133.

27. Kravitz, A.V., Freeze, B.S., Parker, P.R., Kay, K., Thwin, M.T., Deisseroth, K., and Kreitzer, A.C. (2010). Regulation of parkinsonian motor behaviours by optogenetic control of basal ganglia circuitry. Nature 466, 622.

28. Lee, K.-M., Chang, K.-H., and Roh, J.-K. (1999). Subregions within the supplementary motor area activated at different stages of movement preparation and execution. Neuroimage 9, 117–123.

29. Li, N., Chen, T.-W., Guo, Z.V., Gerfen, C.R., and Svoboda, K. (2015). A motor cortex circuit for motor planning and movement. Nature 519, 51–56.

30. Macar, F., Anton, J.-L., Bonnet, M., and Vidal, F. (2004). Timing functions of the supplementary motor area: an event-related fMRI study. Cognitive Brain Research 21, 206–215.

31. Macar, F., Coull, J., and Vidal, F. (2006). The supplementary motor area in motor and perceptual time processing: fMRI studies. Cognitive processing 7, 89–94.

32. Macar, F., Vidal, F., and Casini, L. (1999). The supplementary motor area in motor and sensory timing: evidence from slow brain potential changes. Experimental brain research 125, 271–280.

33. Mantovani, A., Simpson, H.B., Fallon, B.A., Rossi, S., and Lisanby, S.H. (2010). Randomized sham-controlled trial of repetitive transcranial magnetic stimulation in treatment-resistant obsessive–compulsive disorder. International Journal of Neuropsychopharmacology 13, 217–227.

34. Markowitz, J.E., Gillis, W.F., Beron, C.C., Neufeld, S.Q., Robertson, K., Bhagat, N.D., Peterson, R.E., Peterson, E., Hyun, M., and Linderman, S.W. (2018). The striatum organizes 3D behavior via moment-to-moment action selection. Cell 174, 44-58. e17.

35. Mita, A., Mushiake, H., Shima, K., Matsuzaka, Y., and Tanji, J. (2009). Interval time coding by neurons in the presupplementary and supplementary motor areas. Nature neuroscience 12, 502.

36. Mushiake, H., Inase, M., and Tanji, J. (1990). Selective coding of motor sequence in the supplementary motor area of the monkey cerebral cortex. Experimental brain research 82, 208–210.

37. Oh, S.W., Harris, J.A., Ng, L., Winslow, B., Cain, N., Mihalas, S., Wang, Q., Lau, C., Kuan, L., and Henry, A.M. (2014). A mesoscale connectome of the mouse brain. Nature 508, 207–214.

38. Parker, J.G., Marshall, J.D., Ahanonu, B., Wu, Y.-W., Kim, T.H., Grewe, B.F., Zhang, Y., Li, J.Z., Ding, J.B., and Ehlers, M.D. (2018). Diametric neural ensemble dynamics in parkinsonian and dyskinetic states. Nature 557, 177.

39. Pnevmatikakis, E.A., Soudry, D., Gao, Y., Machado, T.A., Merel, J., Pfau, D., Reardon, T., Mu, Y., Lacefield, C., and Yang, W. (2016). Simultaneous denoising, deconvolution, and demixing of calcium imaging data. Neuron 89, 285–299.

40. Pogorelov, V., Xu, M., Smith, H.R., Buchanan, G.F., and Pittenger, C. (2015). Corticostriatal interactions in the generation of tic-like behaviors after local striatal disinhibition. Experimental neurology 265, 122–128.

41. Roland, P.E., Larsen, B., Lassen, N.A., and Skinhoj, E. (1980). Supplementary motor area and other cortical areas in organization of voluntary movements in man. Journal of neurophysiology 43, 118–136.

42. Romo, R., and Schultz, W. (1992). Role of primate basal ganglia and frontal cortex in the internal generation of movements. Experimental brain research 91, 396–407.

43. Rothwell, P.E., Hayton, S.J., Sun, G.L., Fuccillo, M.V., Lim, B.K., and Malenka, R.C. (2015). Input-and output-specific regulation of serial order performance by corticostriatal circuits. Neuron 88, 345–356.

44. Schmidt, R., Leventhal, D.K., Mallet, N., Chen, F., and Berke, J.D. (2013). Canceling actions involves a race between basal ganglia pathways. Nature neuroscience 16, 1118–1124.

45. Stamatakis, A., Schachter, M., Gulati, S., Malanowski, S., Tajik, A., Fritz, C., Trulson, M., and Otte, S. (2018). Simultaneous optogenetics and cellular resolution calcium imaging during active behavior using a miniaturized microscope. Frontiers in systems neuroscience 12, 496.

46. Tarsy, D., Pycock, C., Meldrum, B., and Marsden, C. (1978). Focal contralateral myoclonus produced by inhibition of GABA action in the caudate nucleus of rats. Brain 101, 143–162.

47. Tecuapetla, F., Jin, X., Lima, S.Q., and Costa, R.M. (2016). Complementary contributions of striatal projection pathways to action initiation and execution. Cell 166, 703–715.

48. Viswanathan, A., Jimenez-Shahed, J., Carvallo, J.F.B., and Jankovic, J. (2012). Deep brain stimulation for Tourette syndrome: target selection. Stereotactic functional neurosurgery 90, 213–224.

49. Welter, M.-L., Grabli, D., and Vidailhet, M. (2010). Deep brain stimulation for hyperkinetics disorders: dystonia, tardive dyskinesia, and tics. Current opinion in neurology 23, 420–425.

50. Worbe, Y., Baup, N., Grabli, D., Chaigneau, M., Mounayar, S., McCairn, K., Féger, J., and Tremblay, L. (2008). Behavioral and movement disorders induced by local inhibitory dysfunction in primate striatum. Cerebral cortex 19, 1844–1856.

51. Xu, D., Chen, Y., Delgado, A.M., Hughes, N.C., Zhang, L., Dong, M., and O’Connor, D.H. (2019). A functional cortical network for sensorimotor sequence generation. bioRxiv, 783050.

52. Yin, H.H., Knowlton, B.J., and Balleine, B.W. (2004). Lesions of dorsolateral striatum preserve outcome expectancy but disrupt habit formation in instrumental learning. European journal of neuroscience 19, 181–189.

53. Yin, H.H., Ostlund, S.B., Knowlton, B.J., and Balleine, B.W. (2005). The role of the dorsomedial striatum in instrumental conditioning. European Journal of Neuroscience 22, 513–523.

54. Zhou, P., Resendez, S.L., Rodriguez-Romaguera, J., Jimenez, J.C., Neufeld, S.Q., Giovannucci, A., Friedrich, J., Pnevmatikakis, E.A., Stuber, G.D., and Hen, R. (2018). Efficient and accurate extraction of in vivo calcium signals from microendoscopic video data. Elife 7, e28728.

